# Collagen-based scaffolds with infused anti-VEGF release system as potential cornea substitute for high-risk keratoplasty: An in vitro evaluation

**DOI:** 10.1101/862607

**Authors:** Oleksiy Buznyk, Mohammad Azharuddin, Mohammad M. Islam, Per Fagerholm, Nataliya Pasyechnikova, Hirak K Patra

## Abstract

Currently the only widely accepted corneal blindness treatment is to replace it by transplantation with human donor cornea. However, increasing shortage of donor corneas as well as high risk of rejection in some corneal diseases remain two major problems, which limit the success of corneal transplantation. Corneal neovascularization is considered as one of the main risk factors of graft failure. Different cell-free biosynthetic scaffolds fabricated from collagens or collagen-like peptides are being tested as donor cornea substitutes (DCS). Here, we report for the first-time composite biosynthetic DCS with integrated sustained release system of anti-VEGF drug, bevacizumab and their in vitro validation. We have tethered gold nanoparticles with bevacizumab and integrated into a collagen-based cell-free hydrogel scaffold. Developed grafts preserved good optical properties and were confirmed not toxic to human corneal epithelial cells. Bevacizumab has been shown to constantly releasing from the DCS up to 3 weeks and preserved its anti-angiogenic properties. These results provide background for further use of infused composite biosynthetic DCS with integrated nanosystem of bevacizumab sustained release in corneal disease accompanied by neovascularisation where conventional corneal transplantation might fail.

## Introduction

Corneal diseases are a major cause of vision loss second to cataracts only in overall importance and affect more than 10 million individuals worldwide^1,2^. Currently the only widely accepted corneal blindness treatment is transplantation of human donor cornea. The worldwide demand for transplantation corneas exceeds the supply and this situation will worsen with an aging population and increased use of cornea laser surgery^3^. An alternative for corneal transplantation is replacement of the damaged cornea with an artificial substitute - keratoprosthesis. Existing prostheses neither integrate seamlessly into the host tissue, nor promote reinnervation and mainly remains as a last source option^4^.

Phase I human studies performed by Griffith and Fagerholm^5,6^ in Sweden have shown proof-of-concept for the use of biomimetic materials as corneal implants to promote corneal regeneration as an alternative to donor cornea implantation. Implants fabricated from carbodiimide cross-linked collagen were used in this study as corneal substitute for transplantation and stimulated both corneal tissue and nerve regeneration in the patients, without need for immunosuppression. More recently, we have developed more robust formulations, which were already successfully tested in patients with high risk of donor cornea rejection^7^ and showed that drugs incorporated into nanoparticles, can be released from these corneal implants^8,9^.

This work aims to conquer two big current problems of corneal transplantation. First, developed bioengineered corneal implants can help decrease shortage of human donor corneas, which varies from 30% in developed countries to 80% in developing nations^10,11^. Even if available and transplanted human donor cornea carries a risk of rejection. Major factors responsible for rejection are significant corneal vessels, previous corneal graft rejection in the operated eye, repeated keratoplasty and previous herpetic eye disease^12, 13^. Current ‘gold’ standard of preventing and treating corneal allograft rejection as well as corneal neovascularization (CN) is topical use of corticosteroids. They have well-known side effects such as increased intraocular pressure and cataract formation. Moreover, effect of corticosteroids on CN is not always sufficient. A number of other agents were also tried to inhibit CN such as general and topical cyclosporine, methotrexate and several others with limited effect and side effects^14^.

Excessive VEGF production along with pro-inflammatory cytokines (e.g. transforming growth factors α and ß1 and fibroblast growth factor) are major driving factors for CN. A number of monoclonal antibodies and their derivatives (bevacizumab, ranibizumab, aflibercept) against vascular endothelial growth factor (VEGF) are currently in wide clinical use for inhibition of vessel ingrowth in age-related macular degeneration^15^. They have been also tested as agents to inhibit vessel ingrowth in cornea both before and after corneal transplantation to treat CN and prevent graft failure^14^. However, due to short half-life of the drugs, to the fact that the required dose, the frequency and repetition of the injection are not still defined, it often leads to only partial reduction of CN and its recurrence. Moreover, response in central CN was less marked, probably due to the lower expression of VEGF in long standing vessels and to the distance from the site of injections with respect to a peripheral CN^14^. It was also shown that bevacizumab suppresses CN more effectively than its derivative ranibizumab^16^.

These facts show the need of new approach to treat corneal blindness where conventional corneal transplantation carries a high risk of rejection. Creating a composite corneal substitute with integrated sustained release system of anti-angiogenic drug might become a valuable alternative to human donor cornea allograft combined with frequent anti-angiogenic drug injections or drops.

Objective of the project was to develop composite corneal implants that will serve a dual purpose of: 1) promoting corneal regeneration as an alternative to allograft transplantation, thereby alleviating the organ shortage problem, and 2) delivering anti-vascular endothelial growth factor (anti-VEGF) to stop vessel ingrowth inside the implant, thus decreasing the risk of implant rejection.

## Materials and methods

Bevacizumab (Avastin, La Roche, Switzerland) was used as an anti-VEGF agent throughout the study. All chemicals were purchased from Sigma Aldrich (USA) if not otherwise indicated in the text.

### Synthesis of Gold nanoparticles (GNPs)@Bevacizumab

#### Synthesis of core GNPs

The core GNPs have been synthesized using modified wet-chemical synthesis route as described earlier by Patra et al^17^. Before synthesis of, all the glassware is cleaned with aqua regia (3 HCl:1 HNO_3_) and autoclaved for metallic and non-metallic decontamination. An aqueous solution of 250 μM chloroauric acid (hydrated HAuCl_4_) was brought nearly to the boiling temp at 100°C and stirred continuously with a cleaned and metal ion free stirrer. For the reduction, 600 μM freshly prepared trisodium citrate solution was added quickly at once, resulting in a change in reaction mixture color from pale yellow to deep red to orange in the case of smaller size nanoparticles. After generating the persistent color, temperature was brought down to 25°C, and colloidal GNPs solution was stirred for an additional 15 minutes. After stabilizing overnight, the solution was centrifuged stepwise at 5000 rpm to gently precipitate and re-suspend to make a concentrated stock of 1250 μM GNPs. The particles are then characterized by UV visible spectroscopy and Photon Correlation Spectroscopy (PCS) for hydrodynamic properties along with surface zeta potentials (ξ).

#### Tethering GNPs@Bevacizumab

Considering the sensitive nature of the depletion, we have avoided covalent linking of bevacizumab to GNPs and functionalized the gold nanoparticles (GNPs@Bevacizumab) using electrostatic interactions. While functionalizing bevacizumab, we have considered the fact that amino acids, proteins and polypeptides have multiple types of interactions with GNPs including with the peptide backbone apart from all the positively charged residues^18,19^. Bevacizumab, being a monoclonal antibody (cogitating the isoelectric pH for IgG class ranging from pH 6 to 8)^20^, we choose to trigger the drug in citrate buffer (with pH 5.5) to acquire the net charge positive. GNPs were pre-activated in the same buffer and kept at ξ >-30mV for maximizing the electrostatic interactions and loading of the bevacizumab on surface of the GNPs. The drug was added gradually and until the surface plasmon of GNPs make a drastic change in colour. According to the plasmonic nature of GNPs, we have found that the range of loading could vary from 1-20 mg/ml with an optimum concentration of 11.11 mg/ml (GNPs@Bevacizumab1). The maximum concentration of Bevacizumab loading can go up to 17.85 mg/ml and assigned as GNPs@Bevacizumab2 with pH adjustment. The release of Bevacizumab from the GNPs could trigger by change in ionic strength of the solution and or pH. Particle size and morphology were determined on a JEOL1230 Transmission Electron Microscope (TEM). For characterization and *in vitro* experiments, we have used GNPs@Bevacizumab1.

### Flocculation test

GNPs and GNPs@Bevacizumab were incubated with different concentrations of NaCl solutions (4, 8, 16, 25 and 50 mM) in a 1:1 volume ratio. All the samples were incubated at room temperature for 10 minutes and the hydrodynamic diameter, zeta potential and UV-visible absorbance were measured.^21^

### Fabrication of collagen scaffold with GNPs@Bevacizumab entrapped

Type I porcine atelocollagen was purchased from Nippon Meat Packers Inc. (Tokyo, Japan). Collagen hydrogel scaffolds were prepared as previously described^5,6^. Briefly, 500 mg of 10 % (w/w) porcine type I acidic atelocollagen solution was buffered with 150 μl of 0.625 M MES buffer in a syringe mixing system, 15 μl of 2 M NaOH was added to adjust pH to 5.0. Then, N-hydroxyl succinimide (NHS) and 1-ethyl-3-(3-dimethylaminopropyl) carbodiimide (EDC) were sequentially added in the syringe mixing system and mixed with the collagen solution at 0°C. NHS:EDC: collagen ratio was 0.35:0.7:1. The final mixture was immediately dispensed into glass plate moulds with 500 μm spacer. The hydrogels were cured at 100% humidity at room temperature for 16h, and then demolded in 1X PBS solution. To incorporate free bevacizumab into collagen hydrogels, 50 μl bevacizumab solution 25 mg/ml in 250 μl 0.625 M MES buffer was added to the hydrogel solution prior to adding of EDC to cure. To incorporate GNPs@Bevacizumab, 250 μl of GNPs@Bevacizumab solution 11.11 mg/ml was added into hydrogel solution before adding of EDC. Prepare in similar manner: (a) control hydrogels containing equal amount of bevacizumab stock solution (hydrogel with free bevacizumab), (b) hydrogels containing bare GNPs (hydrogel with GNPs), (c) hydrogel scaffolds with GNPs@Bevacizumab entrapped and (d) hydrogels without drug and nanoparticles (blank).

### Bevacizumab release from the hydrogel scaffolds and GNPs@Bevacizumab using ELISA

Hydrogels containing either free or GNPs encapsulated bevacizumab were placed in 10 ml phosphate buffer saline at 37°C under continuous mechanical shaking and assayed for amounts of the drug released. On days 1, 3, 5, 7, 10, 14 and 24 the PBS was collected and replenished with fresh media. All time points were analyzed by Bevacizumab ELISA kit (ImmunoGuide^®^, Turkey) for amounts of released drug according to kit instructions.

### Fourier Transformed Infrared Spectrosocpy (FTIR) study for Bevaciumab release from collagen hydrogel scaffold

Surface characteristic of bevacizumab released from entrapped hydrogel GNPs@Bevacizumab scaffold from day 1 to 24 were determined by Attenuated Total Reflectance (ATR-FTIR). The measurements were performed using a PIKE MIRacle ATR accessory with a diamond prism in a vertex 70 Spectrometer (Bruker, Massachusetts, USA) with a DLaTGS detector. The whole system was continuously purged with nitrogen and the IR spectra were acquired at 4 cm^-1^ resolution. A total of 64 scans were performed between 4400-600 cm^-1^.

### Cell proliferation assay

Human corneal epithelial cells (HCECs) were purchased from Life Technologies Ltd (Paisley, UK). WST-1 cell proliferation assay reagent was purchased from Roche Diagnostics GmbH (Mannheim, Germany). Keratinocyte serum free medium (KSFM) with supplements-bovine pituitary extract and recombinant EGF (BPE-rEGF), were procured from Life Technologies Ltd.

To assess whether fabricated collagen-based hydrogel scaffolds with Bevacizumab release system inhibit cell proliferation, 1 × 10^4^ cells were seeded on the top of 6 mm hydrogel disc with incorporated GNPs@Bevacizumab in a 96-well plate and cultured for 48h in BPE-rEGF supplemented KSFM at 37°C in 5% CO2. Blank scaffolds, hydrogels with free drug were also cultured along with HCECs culture without any hydrogel scaffold, which served as control for the experiment. In 48h, the media was aspirated and WST-1 reagent was added to the medium following the manufacturer’s instructions. The absorbance was taken at 450 nm in Victor3 V 1420 Multilabel Plate Counter (PerkinElmer, Waltham, MA, USA). Results represent mean values from triplicate measurements.

### Statistical analysis

Difference between groups in Bevacizumab release study and in WST-1 assay was analyzed by Kruskal-Wallis test. A p value<0.05 was considered statistically significant. All the tests were performed on IBM SPSS Statistics 26 and GraphPad Prism 8.

## Results

### Tethering of Bevacizumab on gold nanoparticles and nanosystem characterization

Bioconjugation of bevacizumab on to GNPs surface was studied by DLS, absorption spectroscopy and TEM as shown in Figure 1a-c. Loading efficiency of bevacizumab on to the surface of GNPs was in the range 1-20 mg/ml, we have fabricated two different loading concentration, 11.11 mg/ml of bevacizumab (GNPs@Bevacizumab1) is the optimum concentration of the drug, which can be effectively tethered on to GNPs. The highest concentration is achievable up to 17.85 mg/ml (GNPs@Bevacizumab2). Figure 1(a) represents DLS profile of bare GNPs, GNPs@Bevacizumab1 and GNPs@Bevacizumab2, surface modified GNPs are in the size range of 20 nm in hydrodynamic diameter. This was further verified by UV visible spectroscopy with absorption maxima at 520, 528 and 535 nm respectively for the synthesized nanomaterials (Figure 1 (b)). TEM confirmed nanorange of fabricated nanoparticles with sizes between 15-20 nm.

**Figure 1:**
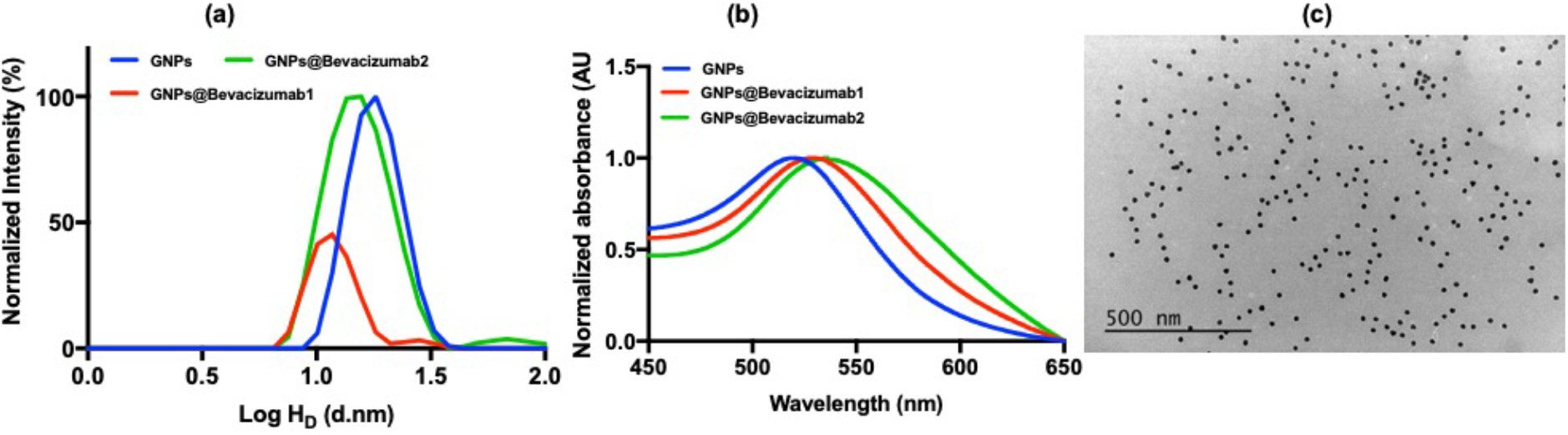
Characterization of bevacizumab tethered gold nanoparticle. (a) DLS of bare GNPs, GNPs surface fabricated with bevacizumab 11.11 mg/ml (GNPs@Bevacizumab1) and 17.85 mg/ml (GNPs@Bevacizumab2), (b) UV-visible absorption spectra for the modified and unmodified GNPs and (c) TEM image of GNPs@Bevacizumab1.

Conjugation of GNPs@Bevacizumab was studied *in vitro* by using salt-induced aggregation study in presence of NaCl as illustrated in Figure 2 a-f. The results depict that the disruption of the electrostatic interaction^22^ between GNPs and bevacizumab leading to the formation of large aggregates (Figure 2(a) & (e)). A higher red shift in the wavelength of the absorption spectra for GNPs@Bevacizumab conjugated species is observed in comparison to the bare GNPs. High concentration of NaCl disrupts the electrical double layer between the particles and induces a shift in the equilibrium between electrostatic repulsion and attraction.^23^ Also, there is an increment in the hydrodynamic diameter of the surface modified GNPs in presence of high salt concentration which further validates to efficient conjugation of GNPs and bevacizumab (Figure 2(e)).

**Figure 2:**
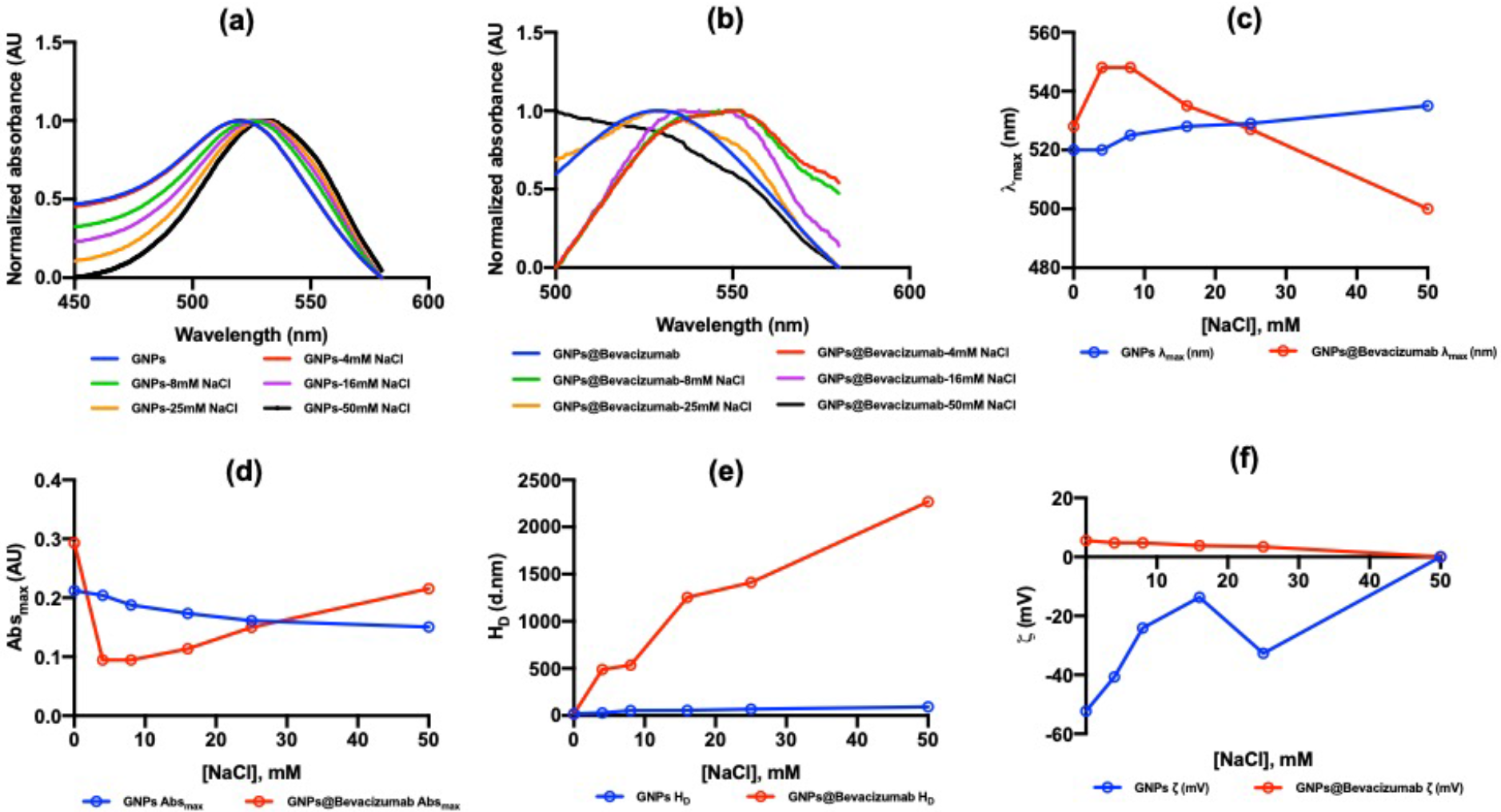
Salt-induced aggregation study (a) Absorption spectral signature of GNPs and GNPs@Bevacizumab (b) in presence of different concentration of NaCl, (c) & (d) lambda max and absorbance value max for GNPs and GNPs@Bevacizumab in presence of varying NaCl concentration. (e) The hydrodynamic diameter and zeta potential of GNPs and GNPs@Bevacizumab in presence of different NaCl concentration.

### Hydrogel with incorporated GNP@Bevacizumab: optical clarity, release profile and cytotoxicity

Incorporation of GNP@Bevacizumab in hydrogel scaffold during fabrication did not influence the scaffold clarity. Composite graft was strong enough to withstand preparation for cell studies i.e. trephination, handling with forceps, which are similar to handling of corneal graft during surgery (Figure 3).

**Figure 3.**
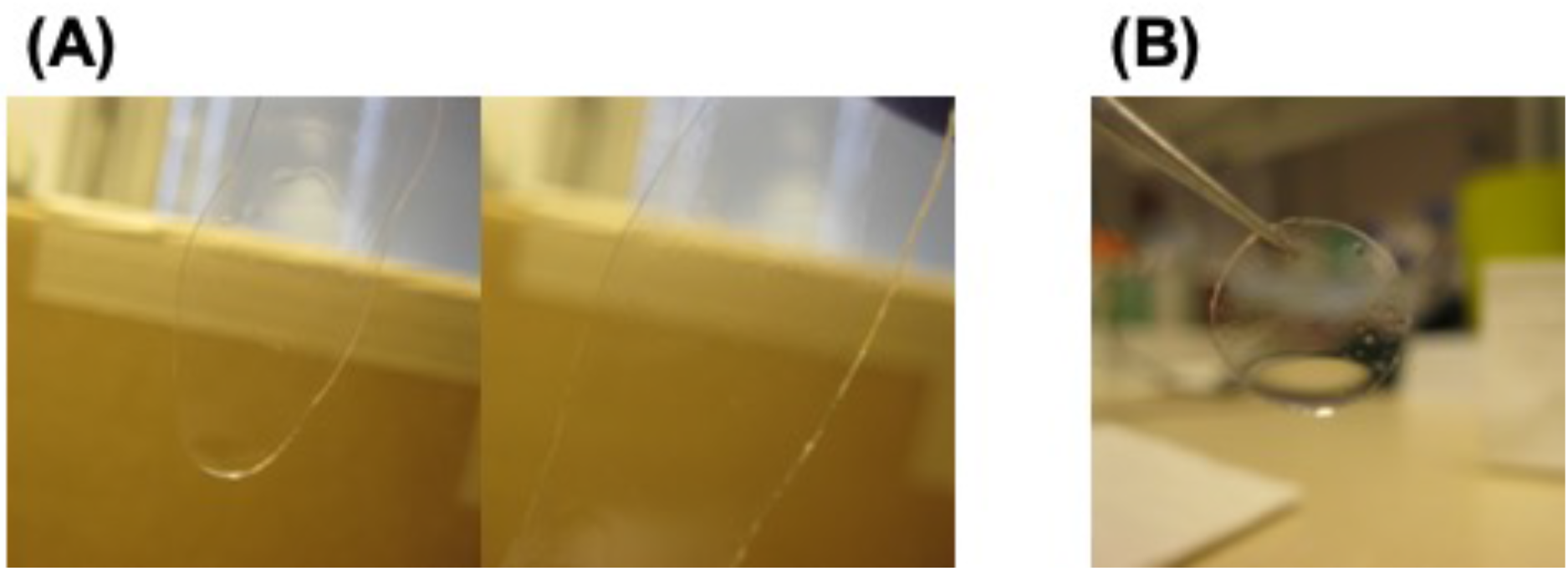
Clear hydrogel scaffold with integrated GNP@Bevacizumab after fabrication (A) and after trephination for cell culture (B).

The sustained release of bevacizumab was obtained with GNP based release system, the drug constantly released up to day 24 both from GNPs not integrated into the hydrogel, and when GNPs@Bevacizumab was incorporated in the hydrogel (Figure 4a-c). The retention time of the drug increased appreciably when it was conjugated with GNPs and integrated into the hydrogel construct as observed by evaluating the concentration of free bevacizumab in the solution using ELISA (p = 0.197). Incorporation of free bevacizumab into collagen implants allowed getting sustained release of the drug up to day 10. 75% of bevacizumab released on day 1 and 90% of free bevacizumab released by day 5. By comparison, GNP encapsulated bevacizumab within collagen implants showed more smooth and gradual release over 14 days. 65% of the drug released on day 1 and 90% of bevacizumab released by day 10 (Figure 4a). Figure 4b represents FTIR spectral signature of bevacizumab released from the hydrogels, the FTIR signature provides two relevant findings: (i) FTIR spectra can be employed for determining the drug release profile of bevacizumab and (ii) conjugation of bevacizumab with GNPs and their entrapment into collagen hydrogel does not show any effect on the secondary structure of the drug (1637 cm^-1^)^24^ as shown in Figure 4 b and c.

**Figure 4:**
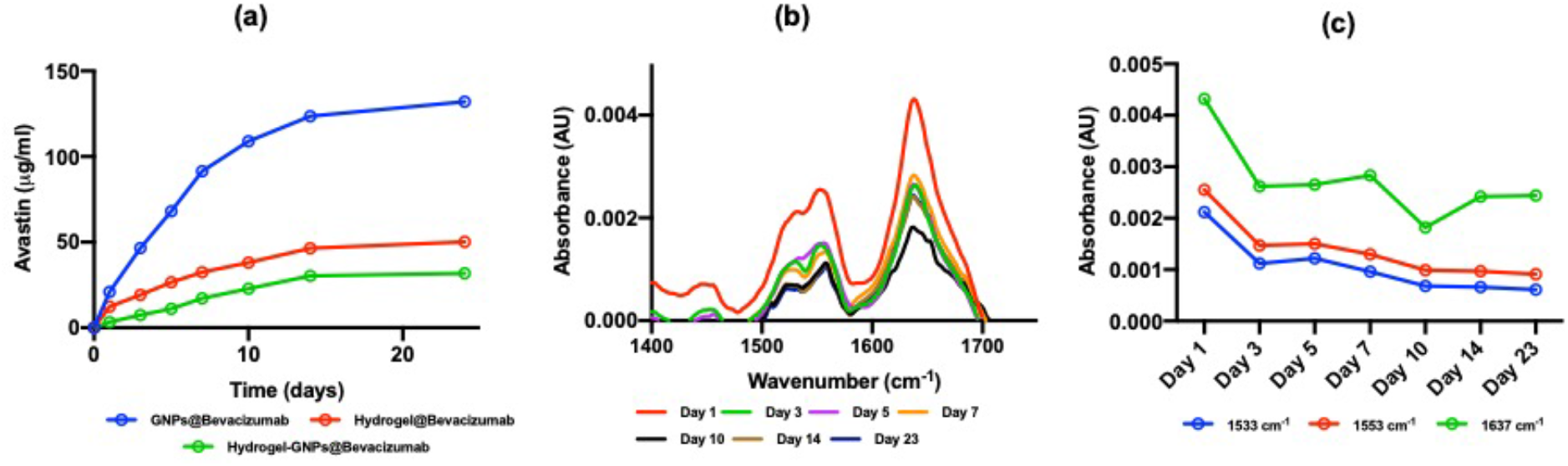
Release profile of bevacizumab from GNPs@Bevacizumab, free bevacizumab entrapped in hydrogel scaffold and GNPs@Bevacizumab entrapped in hydrogel. (a) ELISA for bevacizumab release for the time window (day 1-24) from GNPs@Bevacizumab, free bevacizumab entrapped in collagen hydrogel and GNPs@Bevacizumab entrapped in hydrogel. (b) and (c) FTIR signature of bevacizumab from the released samples day 1-24.

WST-1 cell proliferation assay revealed no cell proliferation inhibition of collagen-based hydrogel with both incorporated free bevacizumab or GNPs@Bevacizumab, when compared to blank/control hydrogel construct and to HCECs cultured on tissue culture plates (p = 0.213). Moreover, drug incorporation into the implants caused light stimulation of HCECs proliferation, but difference was not statistically significant (Figure 5a-e).

**Figure 5:**
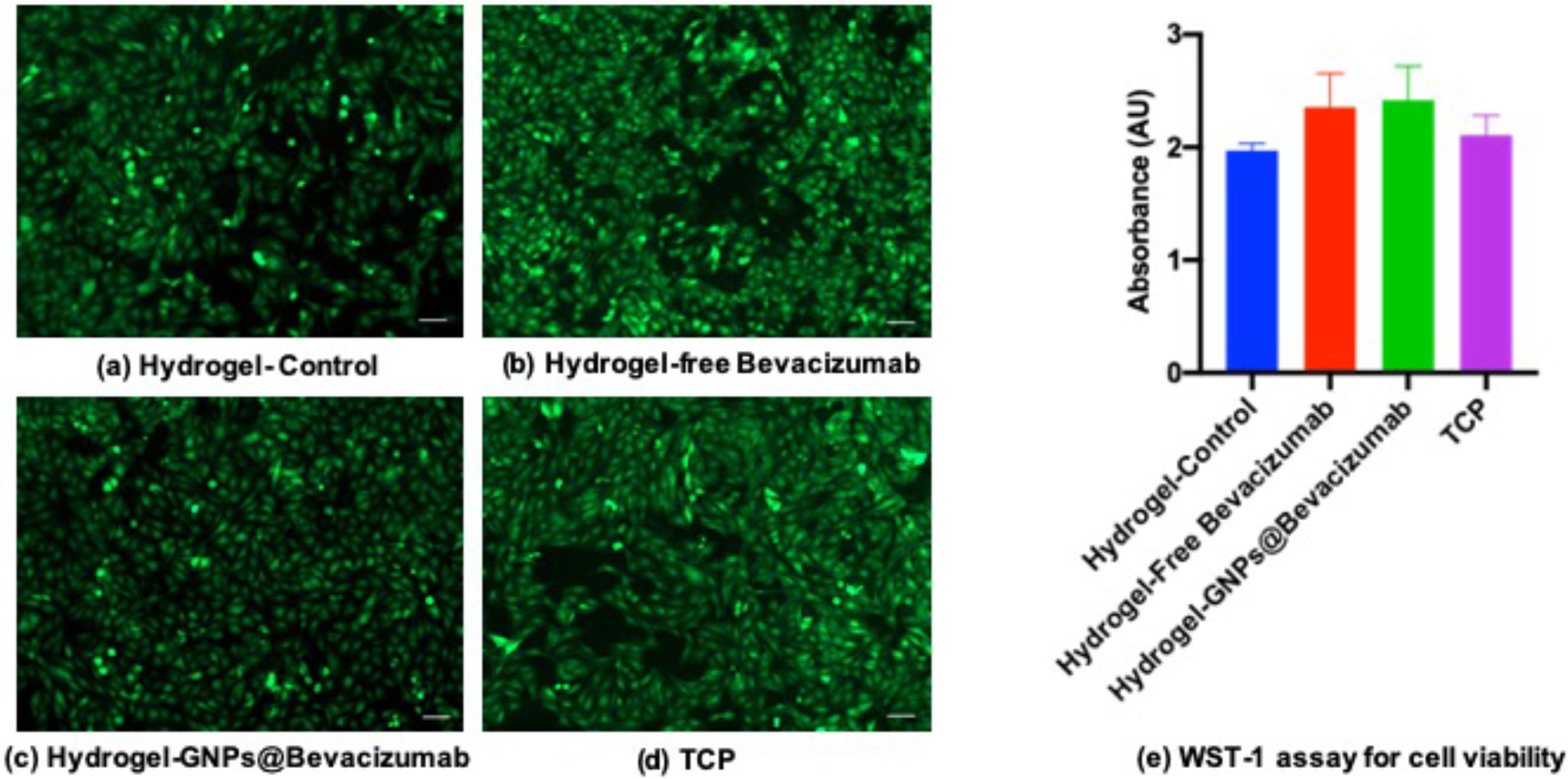
In vitro cell cytotoxicity studies using GFP labelled human corneal epithelial cells for GNPs@Bevacizumab embedded in collagen-based hydrogel scaffold.

## Discussion

Treatment of corneal blindness where conventional corneal transplantation carries a high risk of rejection is of urgent need. More than 50 % of graft rejection has been registered in patients after keratoplasty with preexisting CN^25^. We fabricated for the first time a composite cell-free collagen-based scaffold with integrated sustained release system of anti-angiogenic drug that might become an alternative to human donor cornea allograft combined with frequent anti-angiogenic drug injections or drops in patients with severe corneal disease unsuitable for conventional corneal transplantation or where it has high risk of failure.

Bevacizumab is a monoclonal antibody originally developed for treatment of cancer.^26^ Its ability to inhibit vessel ingrowth due to VEGF blockage has made it one of the most popular drugs in treatment of neovascular age-related macular degeneration.^15^ The main drawback of the drug is that the achieved effect is time limited and it needs to be introduced at least monthly to maintain the effect.^27^ Similar short-term effect of bevacizumab was observed when used for treatment of corneal neovascularization.^28^ That is why fabrication of bevacizumab sustained release system is of great interest and several were already developed and tested *in vitro* and *in vivo*.^29–31^

Bevacizumab attached to gold nanoparticles using electrostatic interactions proved to be a promising option showing sustained release of the drug for up to 24 days. When this sustained release system was introduced in collagen-based corneal implant it did not cause decrease of optical properties of the composite bioengineered corneal graft due to extremely small size of the particles (15-20 nm). Fabricated implants were strong enough to tolerate preparation for cell studies i.e. trephination, handling with forceps, which are similar to handling of corneal graft during surgery. These composite grafts did not inhibit proliferation of human corneal epithelial cells as well. At the same time, sustained release of bevacizumab from collagen-based implant was preserved. More interestingly, incorporation of pure bevacizumab into the implant allowed getting sustained release of the drug as well; still the release profile was not as steep as with GNPs@Bevacizumab.

We suggest that application of developed composite corneal substitute can help alleviating or preventing rejection probability when used in vascularised hazy corneas as corneal allograft due to several reasons. First, collagen-based cell-free hydrogel already has shown low immunogenicity when implanted in both animal and human corneas; no systemic immunosuppression was needed even when they were implanted in high-risk cases.^6–8^ Second, sustained release of bevacizumab from transplanted hydrogel can potentially overcome current problems of the drug use in CN, i.e. short half-life of the drug, not defined required dose, the frequency of the injection. Moreover, this gold nanoparticle-based sustained release system of bevacizumab might be a promising tool for treatment of corneal and retinal neovascularisation when it is used for subconjunctival or intravitreal injections and might decrease need for frequent re-injection of the drug to maintain its anti-angiogenic effect. These findings need further *in vitro* and *in vivo* testing to confirm the suggestions.

In conclusion, we have successfully designed a collagen-based cell-free hydrogel construct with incorporated bevacizumab sustained release system. These constructs are optically clear, strong enough to tolerate surgical handling, non-toxic to corneal epithelial cells and does not influence bevacizumab properties, thus suggesting their suitability, as corneal substitute for application in hazy corneas with neovascularization or high risk of its development. Further *in vitro and in vivo* animal studies needed to be performed to confirm the feasibility of the said nanoparticle entrapped hydrogel scaffolds and also to confirm their efficacy.

## Acknowledgement

We cordially thank Prof May Griffith from Linkoping University, Sweden currently at University of Montreal, Canada for providing research materials and access to the full laboratory set up. This research was supported by EU H2020 Marie Sklodowska-Curie Individual Fellowship (Grant no: 706694) awarded to HP, MIIC Strategic Postdoc Grant awarded to HP and MIIC Seed Grant awarded to HP at Linkoping University (LiU), Sweden. OB was supported by Ögonfonden grant (Sweden). We thank Peter Påhlson for providing us the necessary support and facilities at LiU. OB, MA and HP contributed equally to this work.

